# Exploiting nectar and blood feeding cues and phagostimulants to optimise Attractive Targeted Sugar Baits against a sand fly vector of leishmaniasis

**DOI:** 10.1101/2025.05.07.652609

**Authors:** Daniele Pereira Castro, Fernando Ariel Genta, Matthew Edward Rogers

## Abstract

**Background:** Leishmaniasis presents a major public health problem for a large number of countries requiring effective integrated management of the vector, sand flies, for sustained control. Such strategies need to be economically and environmentally sustainable and adaptable to the behaviour of local vectors. One such tool is Attractive Targeted Sugar Baits (ATSB) that exploit the necessity of sand flies to acquire sugars between bloodmeals. Here we explored the kinetics and cues for sugar and blood feeding to improve the efficacy of ATSBs against sand flies.

**Methods:** A fluorescent assay was developed to quantify sugarmeals to assess the feeding efficiency of colony-reared female *Lutzomyia longipalpis* sand flies.

**Results:** Sand flies showed a range of preferences for different sugars presented on cotton wool and could be manipulated to deposit them into the crop and/or midgut. We found that the combination of 10% sucrose and 10% fructose allowed flies to obtain the largest sugarmeals taken to the crop. Sugarmeals were taken to both the crop and midgut when it contained 200 mM bovine serum albumin (BSA) as a source of protein and 1 mM adenosine triphosphate (ATP) as a phagostimulant. Using this combination, the efficacy of the ingested insecticide fipronil was significantly increased; reducing the 50% lethal concentration from 584 µM to 1.65 µM in a sugarmeal that promoted the simultaneous uptake of the insecticide into the midgut as well as the crop.

**Conclusions:** In this study we highlight the potential of understanding the cues used by vectors to sugar feed and blood feed. By incorporating blood feeding phagostimulants, such as BSA and ATP, in ATSB we vastly improve their killing efficiency against sand flies. This demonstrates a new approach to target these disease vectors.

**Author Summary:** Many insect vectors of disease use sugars, such as nectar, as fuel to enable them to obtain blood, that in turn sustains parasite transmission. In most blood feeding Diptera sugars are diverted to the crop where it is stored and released into the midgut for energy whereas blood is taken directly to the midgut where it is digested to provide nutrients for egg development. For sand flies the cues required to switch between sugar and blood feeding programmes are unknown but if they were they could be exploited to improve Attractive Targeted Sugar Baits (ATSB) as a means of poisoning sand flies when they sugar feed. In this study we find that the sand fly, *Lutzomyia longipalpis*, uses a combination of physical cues (biting/piercing vs sucking) and meal composition to allow them to efficiently feed on nectar and blood. We show that they sense protein and adenosine triphosphate (ATP) to direct blood to the midgut and we demonstrate that this can be turned to our advantage to improve the lethality of ATSB. We show that inclusion of bovine serum albumin as a source of protein and the phagostimulant ATP can trick sand flies to blood feed on sugar and direct the insecticide fipronil to the midgut where it was 355-fold more potent than in sugar alone. These results show the potential of phagostimulants to improve the efficacy and selectivity of ATSB towards blood feeding arthropods, including sand flies, and highlights the value of understanding the feeding programmes of disease vectors in general.

## Introduction

Attractive Targeted Sugar Baits (ATSB) have been shown to effectively control *Anopheles* vectors by providing an ‘attract-and-kill’ approach [1]. They take advantage of the necessity of mosquitoes and other dipteran vectors to feed on sugars for daily survival. It consists of a bait that is attractive to the target vector species, combined with an oral toxin and sugar to stimulate feeding. The toxin is then diverted in to the midgut to exert its effect when the vector requires the sugars for energy to aid their survival, fecundity, flight ability and host-seeking activity. Reducing the vector density is the primary aim of ATSB [2]. With the development of insecticide resistance in malaria vectors and increased outdoor transmission, primary control methods such as insecticide-treated nets (ITN) and indoor residual spraying (IRS) are becoming less effective [3-5]. Therefore, supplementary vector control tools like ATSBs are urgently required. However, to be effective any ATSB toxin needs to access the midgut as quickly as possible and in sufficient amounts to kill the insect.

Female sand flies are the principal vector of leishmaniasis, a group of neglected tropical diseases afflicting people in 99 countries globally with at least 700,000 new cases reported annually [6]. Sand flies acquire and transmit *Leishmania* parasites when they bloodfeed. Between bloodmeals sand flies require sugars, obtaining them from various sources, including, flower nectaries, plant sap, fallen fruits and honeydew from other sap-sucking insects [7-9]. Sand flies use these sugars to fuel their metabolically demanding lifestyle and flight. Sand flies are small (1-3 mm) but can rapidly engorge on blood and take large meals 2-3 times their body size in 1-2 minutes to obtain sufficient protein to produce 30-50 eggs [10,11]. Blood is diverted through the stomodeal valve into the midgut where it is concentrated, surrounded with a peritrophic matrix and digested [12]. Less is known about the sugar feeding habit of these vectors (due to their cryptic nature and distribution across a wide diversity of ecotypes) but they are expected to take small, intermittent nectar meals that are initially deposited into the crop and delivered to the midgut to provide energy as required [13,14]. Therefore, sand flies have distinct blood and nectar feeding programmes allowing them to direct these resources into different organs to be used for different metabolic functions.

Although not known, these programmes are expected to be controlled by sensory input from the meals, cues from the host and the physical act of biting. In this study we sought to begin to uncover these cues by using a fluorescent dye to quantify the volume of sugars imbibed by female *Lu. longipalpis* sand flies and its diversion to the crop and midgut. This allowed us to show that these sand flies mostly use physical cues to activate their bloodfeeding programme but blood-derived phagostimulants could override them to direct sugars into the midgut. We also show that this can be exploited to improve the efficacy of ATSB towards sand flies.

## Materials and Methods

### Reagents

All reagents were sourced from Sigma Aldrich, UK, unless stated otherwise.

### Sand flies

The *Lutzomyia longipalpis* colony of sand flies (originating from Jacobina, Bahia State, Brazil) was maintained at the London School of Hygiene and Tropical Medicine as previously described [18]. Flies were kept at 26 °C and 75% ± 5% relative humidity in darkness. The colony was maintained on fresh human blood and the larvae reared on an autoclaved mixture of compost, daphnia, mouse diet and fine sand. In all experiments unfed female flies 2-3 days post-emergence were used.

### Sugar meal determination assay

Sand flies were exposed to cotton wool pads soaked in various sugar solutions or water containing a fluorescent dye to determine the feeding proportion, the volume of sugar imbibed and the deposition of sugars into the crop and midgut. Female sand flies were transferred to a 15 cm^3^ netting cage and allowed to acclimatise for 2 hours at 26 °C in a humidified (70%) 25 L clear plastic bag. During a feeding experiment, the starved sand flies (3 days post-emergence, deprived access to sugar or water) were supplied with a 6 mm diameter (0.7 g) cotton wool pad (makeup removal pad) placed on top of each cage and soaked with 6 mL 10% sucrose and 10% fructose (w/v, 1:1) dyed with 0.002% (w/v) fluorescein and placed in darkness. After 2 or 4 hours the sugar-soaked pads were removed. Different phagostimulants, 1 mM adenosine triphosphate (ATP),200 mg/mL bovine serum albumin (BSA), and different salts, 140 mM sodium chloride (NaCl) and 25mM sodium bicarbonate (NaHCO_3_), were added to the fluorescein-dyed 10% sucrose-10% fructose solution either singularly or in combination to determine the best sugarmeal components for midgut engorgement.

### Crop and midgut collection and processing

Sand flies were knocked down on ice for 5 mins then washed in a weak detergent solution to remove setae from the body. Washed sand flies were transferred to a dissecting microscope and the entire alimentary tract (crop, midgut, hindgut and Malpighian tubules) was removed as one unit in PBS. To do this the head was removed first using the bevelled edge of a 0.1 mL insulin syringe and the 3^rd^ segment of the abdomen was partially cut. Using the needle the whole alimentary tract was drawn out of the carcass in one careful motion by pulling on the separated tip of the abdomen. The hindgut and Malpighian tubules were trimmed away from the midgut and the crop separated from the midgut. Working quickly, the crop and midgut were pipetted with 5 µL of the dissection fluid into separate 1.5 mL Eppendorf tubes containing 95 µL PBS and placed on ice. Samples were frozen at -40 °C until use.

To measure the fluorescence dissected crops and midguts were freeze-thawed three times by exposing them to hand-hot water for 2 min followed by vortexing for 10 s and dry ice for 2 min. Samples were centrifuged at 2000 rpm for 1 min to pellet any debris before transferring 15 µL to 85 µL PBS in wells of black 96 well flat-bottomed microtitre plates (Nunc). Fluorescence was measured using a Spectramax M3 plate reader (Molecular Devices) set at 485 nm excitation and 520 nm emission with 15 s shaking before reading.

### Crop volume scoring

The sand fly crop was scored for their relative volume. Immediately after dissection the crop was recorded as either empty (presenting as a thin translucent diverticulum, joining the alimentary canal just anterior to the stomodeal valve), partially or fully fed with sugar. Different meals (10% sucrose-10% fructose, blood 50 % in PBS and 50% blood in fluorescein-dyed 10% sucrose-10% fructose solution) were offered to unfed females either through chicken skin, an artificial membrane of parafilm in a Hemotek apparatus or cotton wool in the presence of the human sensory cues CO_2_ and heat (37 °C) for 2 hours.

### Sugar toxin assay

Thirty to forty female unfed sand flies (2-3 days post-emergence) were transferred to 455 mL carboard soup containers (Vegware™), with a 115 mm diameter aperture, covered securely with a fine mesh and allowed to acclimatise for 2 hours at 26 °C in humidified (70%) 5 L boxes. During the toxin-feeding experiment, the starved sand flies were supplied with a cotton wool pad soaked with 6 mL 10% sucrose-10% fructose (w/v, 1:1) with or without 200 mg/ml BSA + 1 mM ATP and varying concentrations of fipronil (1-1000 µM) (RSPCA, UK) or water for 30 min at 26 °C. Sand flies were monitored every 20 min for the first 140 min, followed by a 24 hour endpoint check. Feeding and survival were determined. The number of dead sand flies per total number was used for assessing survival and the median lethal concentration (LC50) was obtained through variable slope nonlinear regression using a Hill equation in GraphPad Prism 10.4.1.

### Statistical Analyses

The p values were determined with GraphPad Prism software, version 10.4.1. As the data was not normal (assessed by a Shapiro–Wilk normality test), the Mann–Whitney unpaired t-test was used to test the value statistical significance between groups for most experiments. All statistical tests were two-tailed.

## Results

### Sand fly sugarmeal intake and fate

Sand flies, like other dipteran vectors, require frequent sugarmeals to maintain their activity which ATSBs exploit as a method for control. To understand the process of sugar uptake we determined the volume of sugar ingested into the *Lu. longipalpis* crop or deposited into the midgut using fluorescein serially diluted and homogenised with an unfed midgut and crop, followed by freeze-thawing. This allowed us to construct a calibration curve and determine that the average crop volume was 12 ± 6 nL (s.d.) after 24 h exposure to our standard rearing concentration of 50% sucrose (1.46 M) (S1 Fig). By comparison the midgut had relative little sugar in it (1.8 ± 2.75 nL). which is consistent with the selective diversion of sugars into the crop. To find an optimal sugar meal for female *Lu. longipalpis* we tested a range of sucrose concentrations and found that 10% resulted in the largest crop volumes (35 ± 18 nL), and the highest proportion of sand flies that fed (31/40, 78%) (Fig. 1A). Next, we tested a panel of mono- and di-saccharides commonly found in nectar [19] at the common concentration of 292 mM. Individually, sucrose resulted in the highest volume imbibed and stored in the crop. In combination with fructose, sand fly crops contained 3- and 4-fold more sugar than either sucrose or fructose alone (Fig 1B), yet the midgut remained relatively devoid of sugar.

**Fig. 1.**
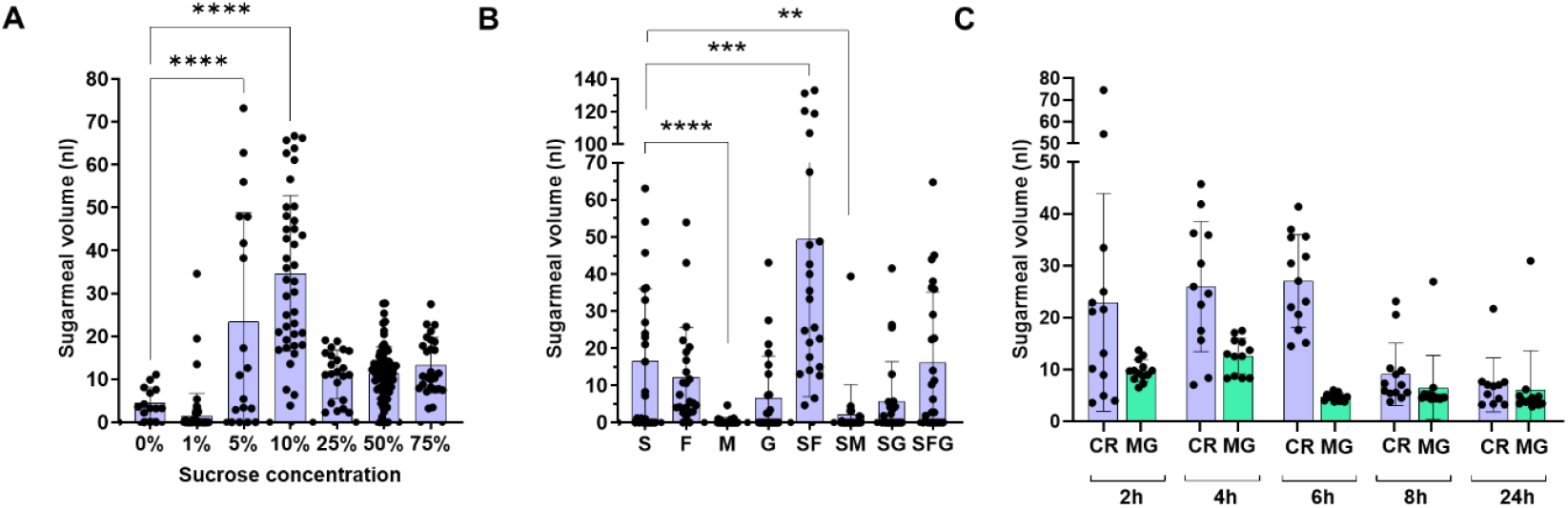
Intake and fate of nectar sugars in female *Lutzomyia longipalpis* sand flies. Unfed sand flies were exposed to different nectar sugars (S = sucrose, F = fructose, M = maltose, G = glucose) dyed with fluorescein. (A and B) Volume of sugarmeals in the crop of sand flies after 4 hours exposure to increasing concentrations of sucrose (A) or different sugars and sugar combinations (10 % final concentration) (B). (C) Volume of 10% sucrose-10% fructose meal in crop (CR) and midgut (MG) over 24 hours after exposure to sugars for 4 hours. Data pooled from 3 independent experiments, n = 10–35/group. Bars represent the mean ± s.d. Asterisks indicate values that are statistically significant (**P ≤ 0.01, ***P ≤ 0.001, ****P ≤ 0.0001) using a two-sided Mann-Whitney unpaired t-test.

To further understand the kinetics of sugar intake and deposition into the midgut we next measured crop and midgut sugar volumes in sand flies exposed to fluorescent meals of 10% sucrose-fructose (292 mM) for 4 hours. Flies were allowed to feed on this sugarmeal and sampled 2, 4, 6, 8 and 24 hours later (Fig 1C). We found that *Lu. longipalpis* load their crop within the 4 hour exposure time and retain this for the following 6 hours during which time the midgut has very little sugar in it, thereafter, the volume of sugar falls in the crop and a consistent small volume can be detected inside the midgut, presumably to be metabolised for energy. These results show that under experimental conditions sugar intake for *Lu. longipalpis* into the crop is quick, stored briefly and then released into the midgut in small volumes over the course of a day.

### Sand flies have separate blood and nectar feeding programmes, but they are not mutually exclusive

Mosquitoes have the ability to discriminate between nectar sugars and blood using specialised neurons within the stylet that pierces the skin [20]. This may allow them to feed regularly on nectar whilst maintaining their appetite for blood. In contrast, the control of the nectar and blood feeding programmes of sand flies is unknown. To see if blood feeding and nectar feeding are separate processes in sand flies, we presented blood and fluorescein-dyed 10% sucrose-fructose to unfed females either through chicken skin, an artificial membrane of parafilm or cotton wool in the presence of the human sensory cues CO_2_ and heat. The proportion of flies that took these meals into their crop or midgut and the relative amount (none, partial, full) was recorded after 2 hours (Fig 2; S2 Fig). As expected, sand flies fed readily on blood though chicken skin in the presence of CO_2_ and heat (100% fed with 87% obtaining full meals in their midgut) and obtain sugars from cotton wool irrespective of the presence of sensory cues (73% fed with 45% obtaining full meals in their crop). A similar result was found for sand flies feeding on blood through parafilm although the overall proportion that fed was much less compared to skin (27% fed with 100% obtaining full meals in their midgut). Flies exposed to blood alone did not take any to the crop, irrespective of the mode of delivery or presence of sensory cues. In contrast, sand flies did not take any blood to the midgut from cotton wool in the presence of CO_2_ and heat but could feed on sugar through skin or an artificial membrane (via skin: 33% fed with 15% obtaining full meals in their crop; via parafilm: 5% fed with none obtaining full meals in their crops). These sugarmeals were found exclusively in the midgut.

**Fig 2.**
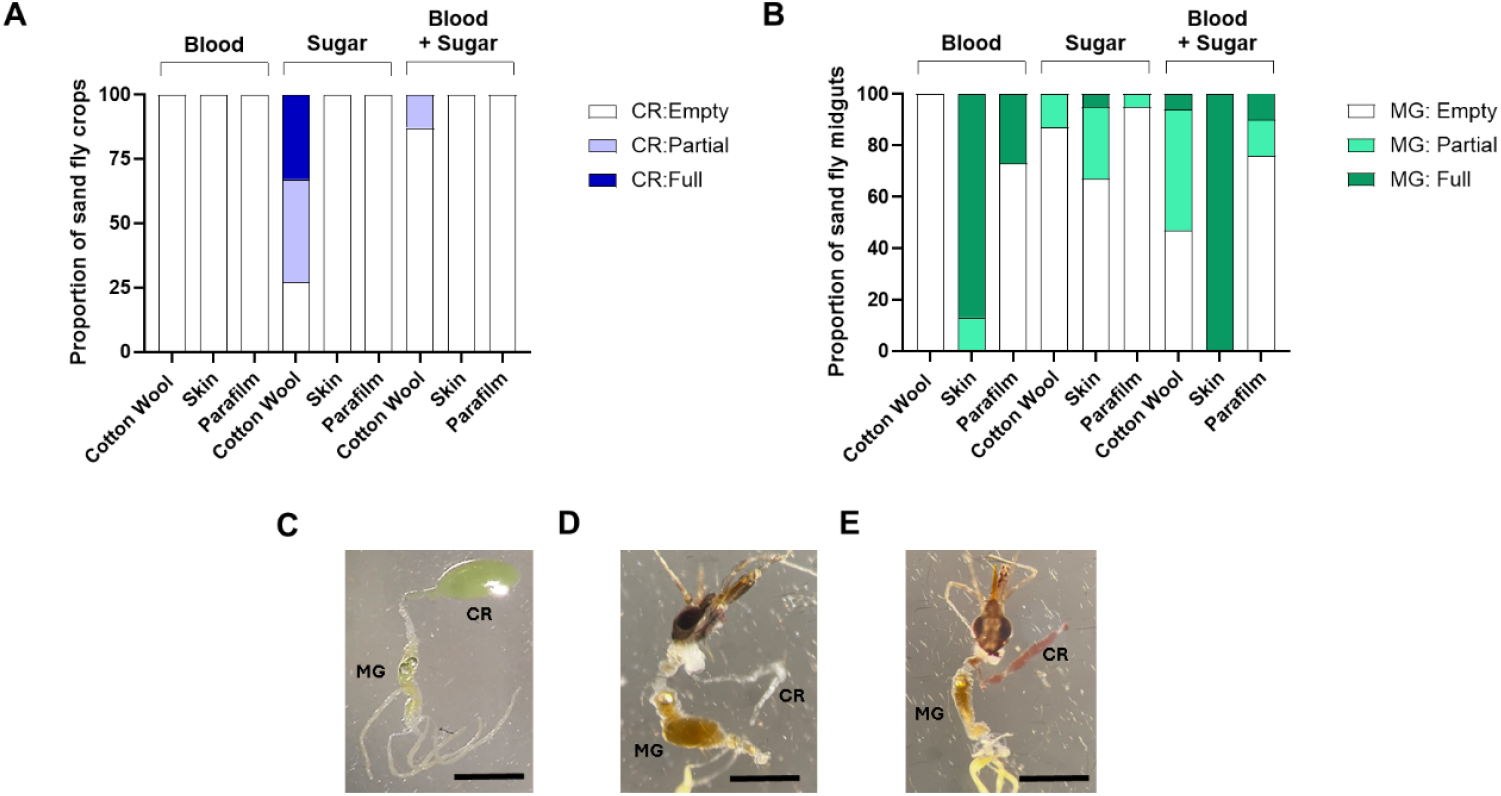
Sugar and blood ingestion through different modes of feeding. Sand flies were fed 10% sucrose-10% fructose, 50 % blood in PBS, or 50% blood in 10 % sucrose-10% fructose solution through cotton wool, chicken skin or parafilm membrane for 2 hours. (A and B) Proportion flies with sugar or blood in the crop (CR) and midgut (MG) and the relative amount (empty, partial meal, full meal). (C-E) Example images of midgut and crop from sand flies fed sugar (C) or blood + sugar (D&E) through cotton wool. Scale bar = 500 µm. Data pooled from 2 independent experiments, n = 30-36/group.

A mixture of blood and sugars resulted in the presence of blood in the midgut and crop of sand flies fed through cotton wool, skin or parafilm (via cotton wool: 53% fed, 100% in the midgut and 25% in the crop; via chicken skin: 100% fed, 100% in the midgut; via parafilm, 34% fed, 100% in the midgut) in the presence of heat and CO_2_. These results highlight the combined effect that the feeding mode (skin penetration, nectar sucking) and meal composition can exert on the feeding programme and destination of the meal. Further, they indicate that although sand flies use different blood and nectar feeding programmes in nature, they are not mutually exclusive and can be potentially manipulated to deliver products to the crop and midgut simultaneously.

### Phagostimulants found in blood divert sugar into the sand fly midgut

From the results of Fig 2 it appears that sand flies can ingest sugar into the midgut but not ingest blood into the crop unless it is offered through a membrane. Previous work with *Aedes aegypti* has shown that it is possible to get these mosquitoes to engorge on an artificial meal of serum proteins or protein-free saline only if the phagostimulant adenosine triphosphate (ATP) was co-present with other plasma components, such as sodium chloride (NaCl) and sodium bicarbonate (NaHCO_3_) [21,22]. This indicates that mosquitoes sense a combination of chemical cues present in blood and nectar to enable them to select an appropriate feeding programme. To see if female sand flies do something similar we tested the if the ingestion of sucrose-fructose into the crop and midgut could be influenced by the presence of the phagostimulants NaCl and NaHCO_3_, bovine serum albumin (BSA) or ATP; either singularly or in combination. Fig 3 shows that after exposure to sucrose-fructose the amount taken to the crop can be substantially increased through the inclusion of BSA + ATP, with a smaller increase shown for flies fed on sugars plus NaCl, NaHCO_3_ + BSA. In all groups the meal volumes in midguts are much lower than crop volumes but between groups the midguts of flies fed sugars with BSA + ATP (with or without NaCl and NaHCO_3_) had more sugar solution diverted to the midgut. Twenty four hours later, crop volumes were lower in the majority of groups and most of the midgut volumes remain unchanged with the notable exception of sand flies fed sugars with BSA+ATP, with or without NaCl and NaHCO_3_. In these flies the average midgut volume increased 16-fold, demonstrating that sand flies could be manipulated to take large sugarmeals and divert them to the midgut.

**Fig 3.**
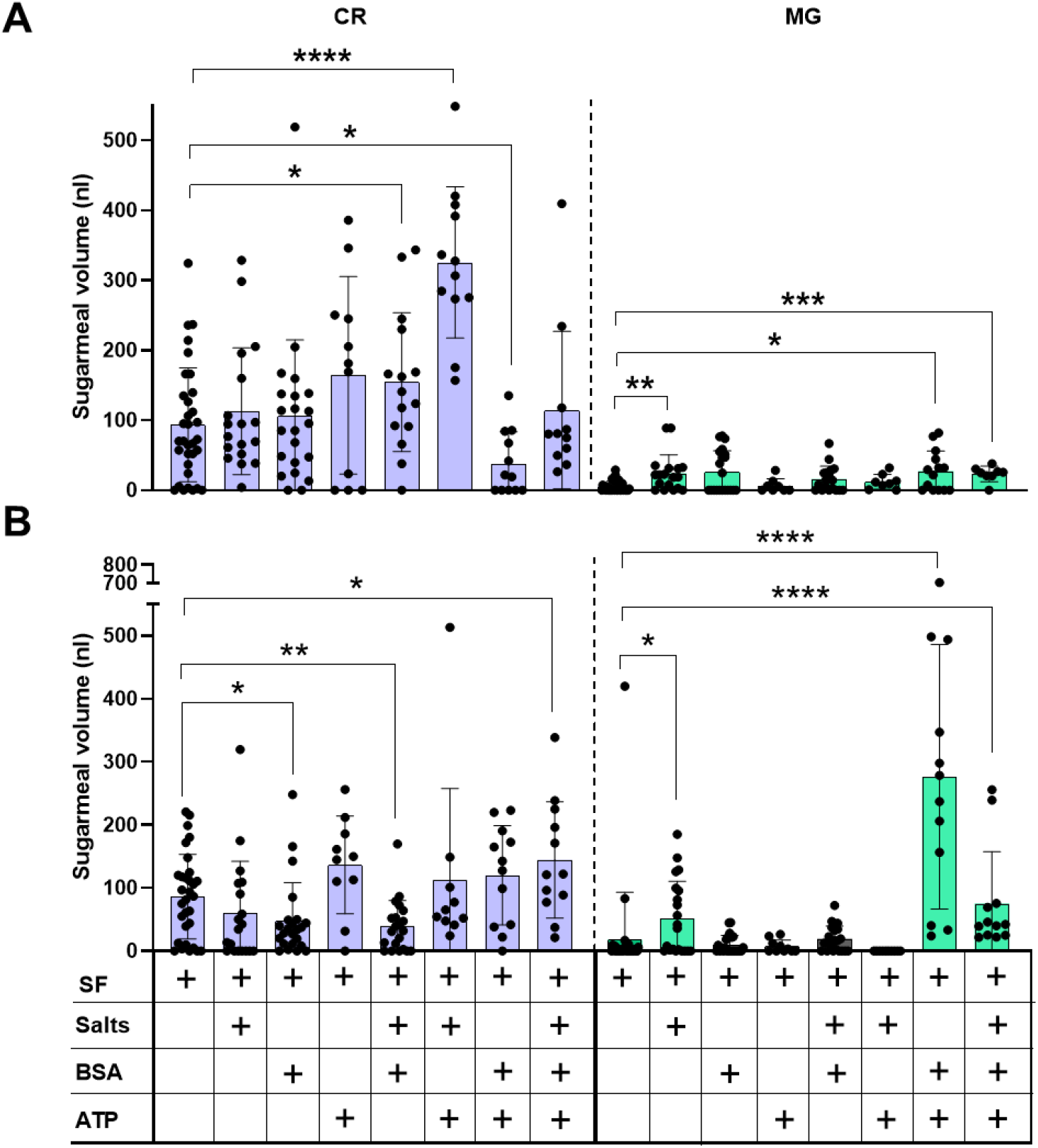
Role of bloodmeal-associated phagostimulants on sugar ingestion and deposition. Volume of meals in crop (CR) and midgut (MG) of female sand flies fed 10 % sucrose-10% fructose (SF) with or without NaCl + NaHCO_3_ (salts), BSA or ATP after 2 hours exposure (A) and 24 hours later (B). Data pooled from 3 independent experiments, n = 10–25/group. Bars represent the mean ± s.d. Asterisks indicate values that are statistically significant (*P ≤ 0.05, **P ≤ 0.01, ***P ≤ 0.001, ****P ≤ 0.0001) using a two-sided Mann-Whitney unpaired t-test.

### Inclusion of phagostimulants in ATSB result in more effective killing of sand flies

Lastly, we tested the potential of mimicking a bloodmeal in an ATSB format for its ability to kill sand flies. Sand flies fed on 10% sucrose-10% fructose with or without the insecticide fipronil (100 µM) showed little difference in the amount ingested and diverted to the crop or midgut (Figs 4A and 4B). However, sand flies fed sugars and insecticide with BSA+ATP took larger meals into the crop and diverted significantly more to the midgut (Fig 4C shows an example image). Over a wide range of concentrations, fipronil in a sugarmeal containing phagostimulants was up to 355 times more potent than in sucrose-fructose alone, with a LC50 of 1.65 µM within by 24 hours of exposure compared to a LC50 of 584 µM for fipronil in sugars alone (Fig 4D and 4E, S3 Fig). This is 8- and 6-fold lower than the topical LC50 and the acute oral LC50 of fipronil to honey bees, respectively [23,24]. The lethality of fipronil in sugar meals with phagostimulants was enhanced and could induce significant mortality in sand flies as early as 60-80 mins post-exposure (Fig 4D). Most concentrations of fipronil tested with phagostimulants (5-1000 µM) showed close to 100% lethality after 24 hours compared to 50% mortality of the group fed sugars and 500-1000 µM insecticide without phagostimulants. Collectively, these results show that the use of phagostimulants to obtain larger sugarmeals that are directed to the midgut substantially improves the lethality of the ATSB towards *Lu. longipalpis* and presents the opportunity to lower the concentration of insecticides or toxicants to reduce non-target effects.

**Fig 4.**
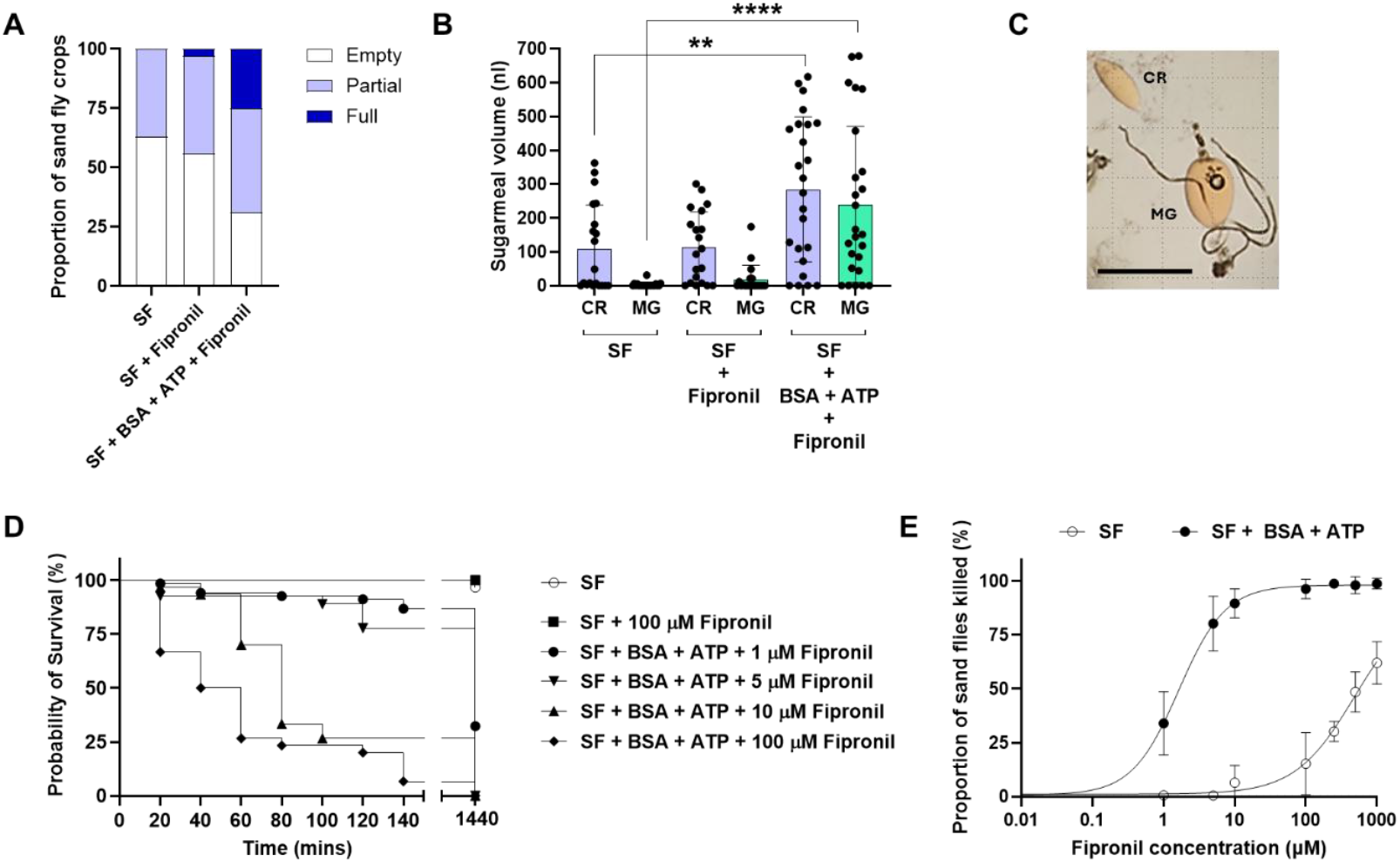
Mock ATSB with phagostimulants against sand flies. Sand flies were exposed to 10% sucrose-10% fructose (SF) with or without phagostimulants (ATP + BSA) or the insecticide fipronil (100 µM). (A) Proportion of crops with a meal of sugar and the relative amount. (B) Volume of sugar in crop (CR) and midgut (MG). (C) Example image of sand fly with a sugarmeal in crop and midgut after feeding on phagostimulants. Scale bar = 500 µm. (D) Kaplan-Meier survival curve of sand flies fed with increasing concentrations of fipronil. (E) Concentration-response curve of fipronil in the presence or absence of phagostimulants in the sugarmeal after 24 hours. Data pooled from 3 (A, B and D) or 4 (E) independent experiments, n = 25-30/group. Bars (B) or symbols (E) represent the mean ± s.d. Asterisks indicate values that are statistically significant (**P ≤ 0.01, ****P ≤ 0.0001) using a two-sided Mann-Whitney unpaired t-test.

## Discussion

Sugar feeding is an essential activity of many dipteran vectors of disease, including sand flies, mosquitoes, black flies, midges and tabanids. Sugars are required to maintain the high metabolic activity of their hematophagous lifestyle, supplying them energy to locate bloodmeals and egg-laying sites in order to use the proteins and lipids from blood to generate large batches of eggs. Thus sugar feeding has the enormous potential to be exploited for the control disease vectors, as a means of introducing toxicants, insecticides, anti-parasitics, anti-feeding agents or microbes expressing inhibitory RNAs into them. However, Attractive Targeted Sugar Baits (ATSB) currently rely on sugars as the sole phagostimulant. This presents a problem as many vector species take small, intermittent sugarmeals from a variety of sources. This means that multiple visits to the bait would be required to achieve toxicity.

In recent years there has been a drive to understand the feeding behaviour of mosquitoes and this has identified a number of phagostimulants that show promise in improving ATSBs [20,25]. In this study we sought to understand the sugar and blood feeding response of sand flies to improve the ingestion of sugar and its diversion to the midgut – the site of action for many insecticides and toxicants. Adult sand flies of both sexes feed routinely in sugar sources as nectar, honeydew, or fruit juices, and only females require a blood meal from a vertebrate host for egg development [26-28]. In laboratory or field conditions, sand fly sugar meals are mimicked artificially using sugar baits (e.g. cotton wool embedded in a sugar solution). Importantly, it has been shown that the ingestion of sugar by the sand fly is an important stimulus for the complete development of the *Leishmania* parasite [29].

To compete in the field with the vector’s natural sugar sources, the quality of the attractant is a crucial component of the ATSB design. Equally important but rarely considered is the quality of the meal itself - to promote as much toxin or insecticide to be taken during a single feed. This may be a missed opportunity as, currently, no ATSB has been designed to get a vector to engorge on a sugarmeal and directly target the insect’s midgut.

To see if this was possible for sand flies we used a robust fluorometric assay to sensitively measure the quantity of a sugar in the crop and midgut of the neotropical leishmaniasis vector, *Lu. longipalpis*, to assess the effect of different sugars baits on meal size and how quickly it can access the midgut. We show that the combination of sucrose and fructose provided the best sugar source, resulting in the largest crop meals within hours of exposure to the sugar source. Interestingly, glucose, the only sugar found in both blood and nectar, was infrequently taken into the crop. This suggests that for sand flies nectar and blood feeding cues are largely distinct but show a degree of overlap that may be sensed, in the context of other cues, to efficiently switch between these two feeding modes. In seminal work by Hosoi and Galun the meal components for the dengue vector *Ae. aegypti* to engorge were determined and, importantly, were shown to work without blood [30-32]. They demonstrated that blood feeding could be uncoupled from its value as a protein-source by showing that ATP induced engorgement only when co-presented with plasma components such as NaCl and NaHCO_3_. Later, Jove et al. (2020) elegantly demonstrated that female *Ae. aegypti* employs specialist neurons in the stylet (the mouthparts that pierces the skin and cannulate blood vessels) to discriminate nectar and blood and are insensitive to the nectar sugars sucrose and fructose [20]. They are activated by glucose only in combination with other bloodmeal cues, including NaCl, NaHCO_3_, ATP and serum proteins. Interestingly, only half of the neurons were activated in response to bloodmeal cues indicating that others are important but are as yet unknown.

In a simple yet revealing, experiment, we showed here that female sand flies ingest blood only when presented through a membrane (of skin or parafilm) and not through cotton wool, unlike sugars that could be consumed from cotton wool or through a membrane. However, a mixture of nectar sugars and blood on cotton wool resulted in a significant proportion of flies with blood in their crop and midgut. This indicates that the destination of a meal in sand flies can be manipulated according to its constituents and that cues in blood have the potential to direct a sugarmeal to the midgut. In mosquitoes nectar is taken to the crop, whereas blood bypasses the crop and is routed to the midgut, where it is digested. Mosquitoes will readily engorge on blood through a membrane provided with host-sensory cues such as heat and CO_2_, however, if the bloodmeal is replaced with sugars the mosquitoes will reject this meal if these sensory cues are present (Bishop and Gilchrist, 1946) [33]. This indicates that cues in the odour and the taste of a meal allow these vectors to flexibly chose which feeding programme to employ. To this end, Zhilin et al. (2022) found that activation of ATP-sensitive potassium channels in the salivary glands misdirected blood into the crop [34]. Collectively, this offers the premise to explore phagostimulants, such as ATP, as a means to improve ATSB function. In contrast, little is known of the mechanisms underlying sugar and blood feeding in sand flies, however, what we do know may be quite revealing. Physical cues appear to be important and how the sand flies use their mouthparts. Meals derived from plant surfaces, such as honeydew from aphids or flower nectar, were observed to be diverted to the crop, whereas meals taken directly from plants through piercing, as seen in blood feeding, are largely directed to the midgut [13]. Using dyed sugarmeals, Tang and Ward showed that female and male *Lu. longipalpis* sand flies interrupted during feeding had traces of sugar in the midgut, but that the rest of the meal was diverted into the crop [35], something that was also seen in this study. Next, they investigated the ultrastructure of the stomodeal valve (an invagination of the midgut with an underlying ring of muscle that allows the midgut to retain ingested blood) finding basiconic sensilla on the inner side of the oesophagus at the junction with the stomodeal valve [36]. They proposed that the sensilla may control the movement of the valve after contact with fluids entering the midgut and may be responsible for diverting the sugarmeal to the crop by closure of the valve only after a small volume of sugar has passed through it. Supporting this, we could detect the fluorescent sugar meal in the midgut of *Lu. longipalpis*, albeit at low levels, as early as 2 hours after exposure to a sucrose-fructose meal.

All blood-sucking insects use a range of phagostimulants to discriminate sugar and blood that could allow them to obtain large reserves of sugar without affecting their appetite for blood. Therefore, we tested the effect of a range of blood feeding cues to encourage female sand flies to ingest large sugarmeals rapidly into their crops and divert it directly to the midgut – i.e. trick them to blood feed on the sugar bait. A mixture of four components of blood - BSA, ATP, NaCl and NaHCO_3_ - were tested. We found that ATP could induce sand flies to take large sugarmeals into their crop, with the largest (548 nL) found in the combination with NaCl and NaHCO_3_. After 24 hours the flies with the largest sugarmeals remaining in the crop were those presented ATP in the sugar solution (in combination with any of the other cues), however, in these flies there was little or no sugar diverted to the midgut. By far the largest diversion of sugar occurred after 24 hours in sand flies fed with a combination of ATP and BSA - capable of introducing 709 nL into the midgut. This compares well with the volume of blood that a fully engorged female *Lu. longipalpis* can take (1 µL [10]) and the mean bloodmeal volume ingested by a number of other phlebotomine sand fly species (0.47 to 1.01 µL [11]). Confirming this result, a follow up experiment found that the combination of BSA and ATP led to the largest proportion of female sand flies with full crops; larger amounts of sugar taken into the crop and diverted to the midgut and dramatically more rapid killing of sand flies using the example insecticide fipronil, commonly used in ATSB (Fig 4).

By introducing cues that promote rapid midgut engorgement on sugars there is scope to improve ATSB to control vector-borne diseases. In our mock ATSB we show the potential of combining phagostimulants with the sugar bait and how they can be improved to be more lethal more quickly against sand flies and, potentially, other disease vectors. Although not tested in this study, lowering the dose of toxicant or insecticide may also substantially reduce their off-target effects on insect species that also nectar feed, such as pollinators and has the potential to improve the effects of ATSB targeting the *Leishmania* parasite [26, 37]. With this in mind, careful combination of blood feeding cues may also improve the selectivity of a ATSB towards blood-sucking arthropods but more needs to be known about them and how vectors other than mosquitoes sense and switch between nectar and blood feeding programmes. Sand flies belong to a subgroup of disease vectors that feed upon a pool of blood by lacerating the upper dermal capillary loops in the skin. Pool feeding may generate unique cues that mosquitoes may not encounter, such as skin constituents or wound-associated cytokines, chemokines and volatiles [38], so it would be prudent to explore these in greater detail to expand the range of vectors of this emerging control tool.

## Acknowledgements

We would like to thank Ms Shahida Begum for assistance with sand fly colony rearing at LSHTM.

## Financial Disclosure Statement

DPC was funded by a CAPES - SENIOR VISITING PROFESSOR CAPES-PRINT - 88887.878914/2023-00. FAG was funded by CNPq – Produtividade em Pesquisa - 312305/2022-2 and FAPERJ – Cientista do Nosso Estado – E-26/200.454/2023. MER was supported by the BBSRC (David Phillips Fellowship awarded to MER, BB/H022406/1). The funders had no role in study design, data collection and analysis, decision to publish, or preparation of the manuscript.

## Notes

### Competing Interest Statement

The authors have declared no competing interest.

