## Supplementary Figures for "Exploiting nectar and blood feeding cues and phagostimulants to optimise Attractive Targeted Sugar Baits against a sand fly vector of leishmaniasis"

**Short title: Phagostimulants improve ATSB efficacy against sand flies.**

### Supporting information

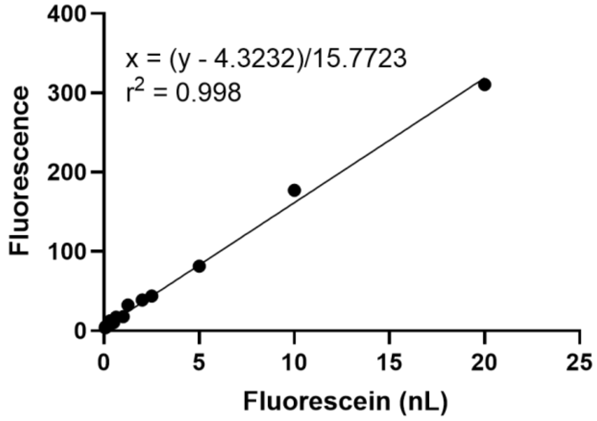


S1 Fig

**S1 Fig. Calibration curve for quantitation of sugarmeal volume in a sand fly crop or midgut.** A serial dilution of 50% sucrose dyed with fluorescein was made with a crop and midgut from a single unfed female sand fly. For each dilution, sugar, dye and dissected organs were homogenised before reading the fluorescence (485 nm excitation and 520 nm emission). Values represent the average of 4 replicates per dilution.

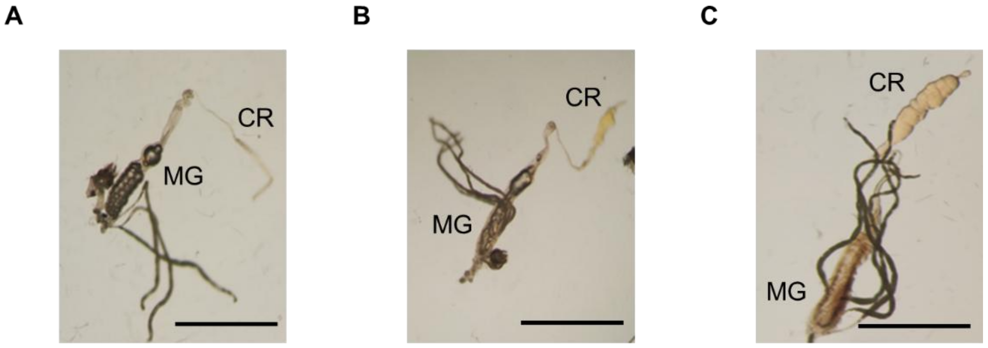


S2 Fig

**S2 Fig. Representative images of sugarfed crops from sand flies.** Flies were fed on 10% sucrose-10% fructose solution dyed with 0.002% fluorescein through cotton wool. (A) empty crop, (B) partial sugarmeal or (C) full sugarmeal. Crop (CR), midgut (MG). Scale bar = 500 µm.

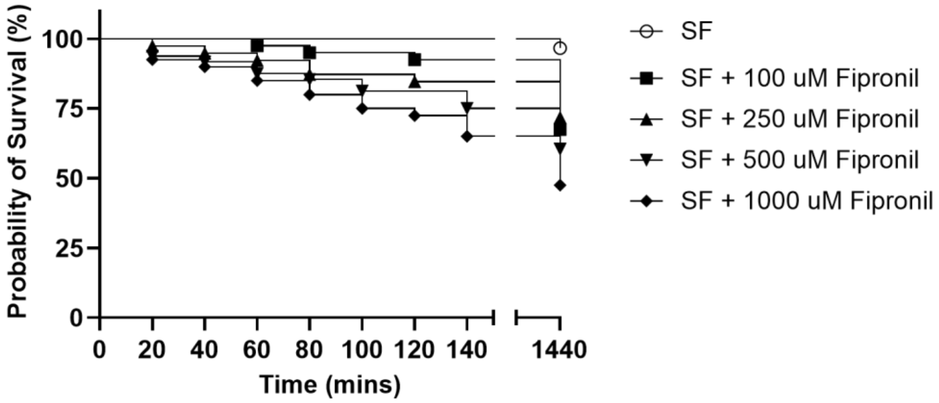


S3 Fig

**S3 Fig. Mock ATSB against sand flies, without phagostimulants.** Kaplan-Meier plot of survival of sand flies fed with increasing concentrations of fipronil. Sand flies were exposed to 10% sucrose-10% fructose with or without the insecticide fipronil (100-1000 µM). Data pooled from 3 independent experiments, n = 39-59/group.
